# Automated identification and registration of anatomical landmarks in *C. elegans*

**DOI:** 10.1101/2022.03.29.486182

**Authors:** Nicolette M. Laird, Zachary Pincus

## Abstract

The physiology of the nematode *C. elegans* can be visualized with many microscopy techniques. However, quantitative microscopy of *C. elegans* is complicated by the flexible and deformable nature of the nematode. These differences in posture and shape must be addressed in some fashion in any automated or manual analysis. Manual approaches are time intensive and require hand-labeling anatomical regions of interest. Automated tools exist, but generally rely on high-magnification imaging using labeled nuclei as fiducial markers. Here we describe a suite of new tools that allows for high-throughput analysis of whole-body images, aligned using anatomical landmarks identified from brightfield images. We show how these tools can be used in basic morphometric tasks and examine anatomical variation and morphological changes in a population over time.

## Introduction

In the last decade, a variety of high-throughput, image-based approaches have been developed to measure phenotypes in *C. elegans*. Systems like the Wormotel (1), Worm Corral (2), and Lifespan Machine (3), produce large amounts of imaging data that must be processed and analyzed. A key challenge with such systems is that the animals are often free-moving and can adopt a variety of positional conformations, making it difficult to compare images quantitatively across individuals. Some microfluidic devices solve this problem physically, with constricted channels into which individuals can be pulled for imaging (4–7). In other cases, such approaches are not feasible, especially for culture of *C. elegans* on solid media. Computational techniques to account for positional and anatomical differences among individuals are thus critical to fully exploit images obtained in these systems.

The first step in computationally analyzing imaging data requires identifying the regions of interest, a process known as “image segmentation”. Recent advances in computer vision have made it easier to find objects in images, but identifying pixels belonging to individual worms in noisy or cluttered images is still a challenge. Traditional image analysis techniques like thresholding, the watershed transform, and edge detection have been effective at segmenting images where there is a strong difference in pixel intensity between the background and the object. These techniques have been used to automatically identify *C. elegans* animals in several high-throughput assays because in many lower magnification imaging conditions, *C. elegans* often appear much darker compared to their backgrounds (8–11). Slightly higher magnifications and/or imaging through media with a refractive index closer to that of *C. elegans* improve resolution of internal structures but often decrease the contrast between animal regions and the background, making segmentation challenging. In such conditions, as arise in our Worm Corral system (See Methods), more sophisticated deep-learning-based approaches show promise in reliably separating individuals from cluttered backgrounds (12).

However, distinguishing worm regions from the background is not sufficient to extract morphological data from images. Due to their flexible and deformable nature, *C. elegans* can adopt a variety of complex postures, which can obscure automated feature detection. Several algorithms have been developed to address this challenge by computationally straightening the curved worm body (8–11,13–16).These algorithms tend to follow similar steps: I) identify the set of image pixels corresponding to an individual *C. elegans* (via segmentation) II) use the identified pixels to determine the centerline of the individual, III) use the centerline to computationally straighten the curved worm body.

Beyond overall posture, a second major challenge in comparing image patterns across individuals is internal anatomical variation. In the past, several tools for analyzing *C. elegans* images at single-cell resolution have used image registration methods to account for variation in the position of cellular structures across individuals and over time (8,9,17–23). The goal of image registration is to bring two images into spatial alignment, often using nonrigid warping transformations (24, 25). Such methods have been employed in *C. elegans* to ensure individual cells are correctly identified in embryos, developing animals, and neuronal studies (8,9,17–23). Many of these methods rely on high-magnification imaging and fluorescent staining or transgenic reporters to define landmarks (also known as fiducial markers) that are then used to align images (8,9,20,26). For lower magnification or whole-worm imaging, however, such techniques have limited applications, especially, in conditions where images of fluorescent landmarks are hard to acquire.

Here we present a suite of fully automated tools to correct both positional and internal variation across adult whole-body *C. elegans* images without the need for fluorescence imaging (Fig 1A). Our method relies on identifying each animal’s centerline from a segmented image, defining curving but locally orthogonal anterior–posterior (AP) and dorsal–ventral (DV) axes (Fig 1B). Given these axes, it is possible to transform original images in the “laboratory frame of reference”, where the x and y coordinates refer the axes of the imaging apparatus, into a “worm frame of reference”, where the animal’s AP and DV axes are warped to align with the image’s axes; this produces an image in which the worm is computationally straightened. By transforming between coordinates in the worm frame and lab frame of reference, positional variation between individuals can be accounted for (Fig 1C). We next automatically identify anatomical landmarks from brightfield images to further bring images into register, addressing variations in internal anatomy across individuals. This allows the straightening and internal alignment of any set of imaging data as long as corresponding brightfield images are available. Finally, we demonstrate how this approach can be used to quantify population trends in anatomical variation in a dataset comprising thousands of images of a 118 individual worms observed from young adulthood until death.

**Fig 1.**
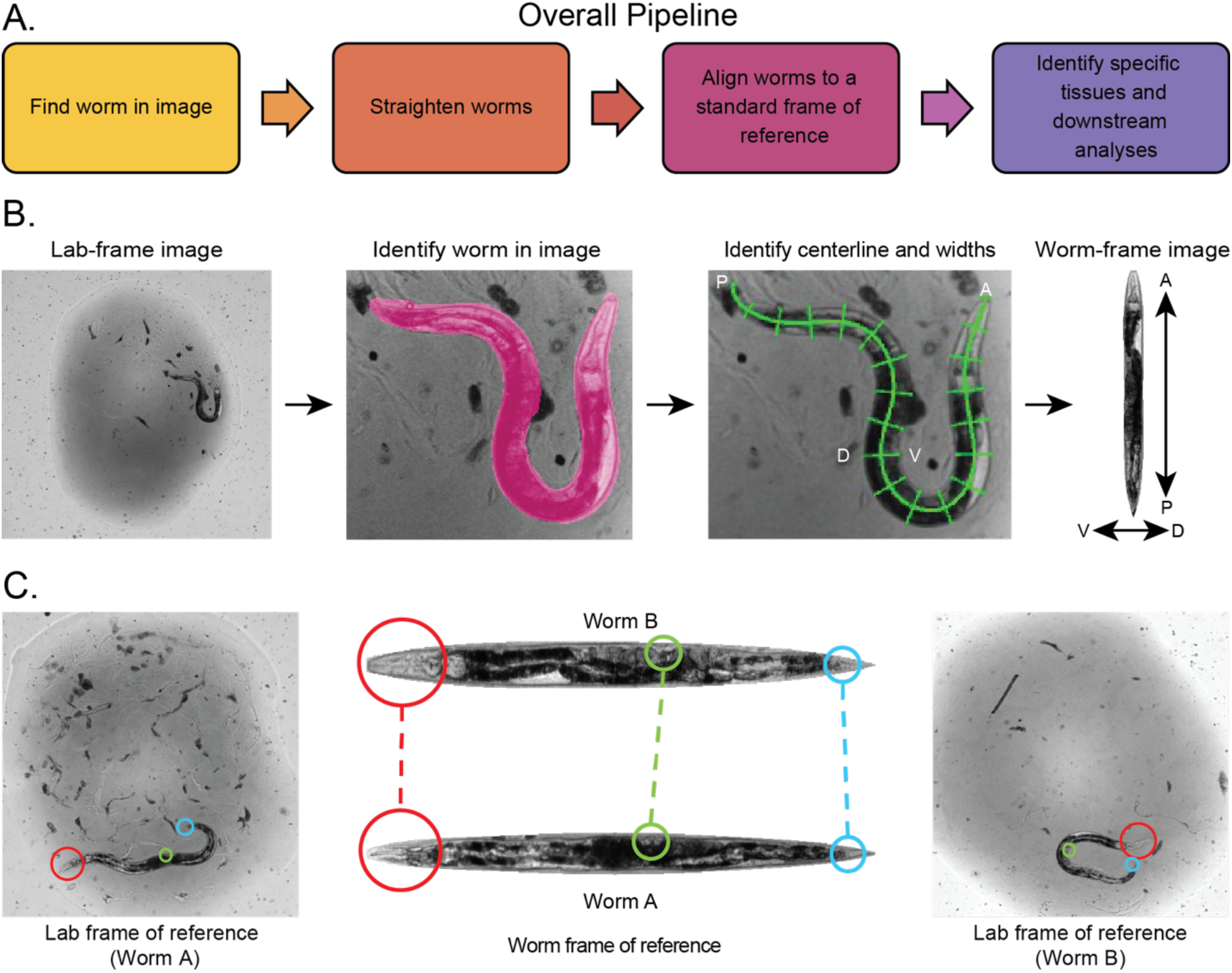
A method to establish point correspondences across *C. elegans* images. (A) Overall pipeline. (B) General method for pose estimation and computational worm straightening. A binary mask of the worm pixels is automatically generated from a brightfield image (the “lab frame of reference”). The nose-to-tail centerline and perpendicular widths of the animal are identified from the mask. These two axes define a new coordinate system local to the individual, the position along its anterior–posterior and dorsal–ventral axes. In particular, these axes allow the (x, y) coordinates in the original (“lab-frame”) image to be determined for any position specified along the animal’s anterior–posterior (AP) and dorsal–ventral (DV) axes. This defines a (nonrigid) coordinate transformation that allows the “lab-frame” image to be resampled along the individual’s AP and DV axes, producing an image in the “worm frame of reference”. (C) The worm frame of reference enables point correspondences to be made between different images. Regions corresponding to specific anatomical locations can be defined in the worm frame of reference and, for each image of interest, mapped back to the original lab frame of reference. Measurements can then be made on the corresponding pixels of the original lab-frame image without additional transformations or could be made directly on the worm-frame image.

## Results

### Accounting for positional variation

To compare image data across individuals and temporally, it is first necessary to account for the effects of the animal’s orientation and posture within each image. This requires identifying regions in separate images that correspond to the anatomical structures of interest. Manual efforts such as circling the regions corresponding to a specific neuron across a set of fluorescent images, for example, are a special case of this general task. A systematic computational approach to the problem is to define a worm-local coordinate system, such that each image pixel within the animal can be labeled with two coordinates: the pixel’s position nose-to-tail along the animal’s anterior–posterior (AP) axis, and in the case of 2D imaging of animals crawling on solid media, its position along the dorsal–ventral (DV) axis, which is locally orthogonal to the AP axis (Fig 1B). Locating an animal’s centerline (see Methods for details) is sufficient for this task, as the centerline defines the AP axis and the vectors normal to every point along the centerline define the DV axis. Once these axes are determined, it is simple to generate a straightened “worm frame of reference” image by sampling image intensities along a grid of anterior–posterior and dorsal–ventral positions (Fig 1B). In this way, whole-animal images can be directly aligned and compared. Alternately, it is also possible to measure localized image intensity patterns without any image transformation by simply defining sub- regions of interest in the AP/DV coordinate system. These “worm-frame” coordinates can be transformed back into pixel regions in any particular “lab-frame” image, allowing raw, untransformed pixel values from those regions to be compared directly (Fig 1C).

Moreover, if a brightfield and fluorescence image are taken in quick enough succession such that the worm has not substantially changed its posture, then the centerline and outline found from the brightfield image can be applied to the fluorescence image as well.

### Straightened “Worm-frame” images remain valid for quantitative comparison

At its core, transforming an image from the lab frame into the worm frame is a sampling problem. Computationally straightening the curved worm body involves mapping points from one coordinate system (the lab frame of reference) to another (the worm frame of reference). A potential problem arises, however, when mapping points along a highly curved shape like that of a coiled animal. Because the arc length differs between the inner and outer edges of a curved shape, sampling image intensities at single-pixel spacing along the centerline produces a relative undersampling of pixel intensities along the outer edge and a relative oversampling of intensities along the inner (Fig 2). This results in image intensities along the outer edges being under-represented in worm-frame images, and those along the inner edges being duplicated. An example is depicted in Fig 2A.

**Fig 2.**
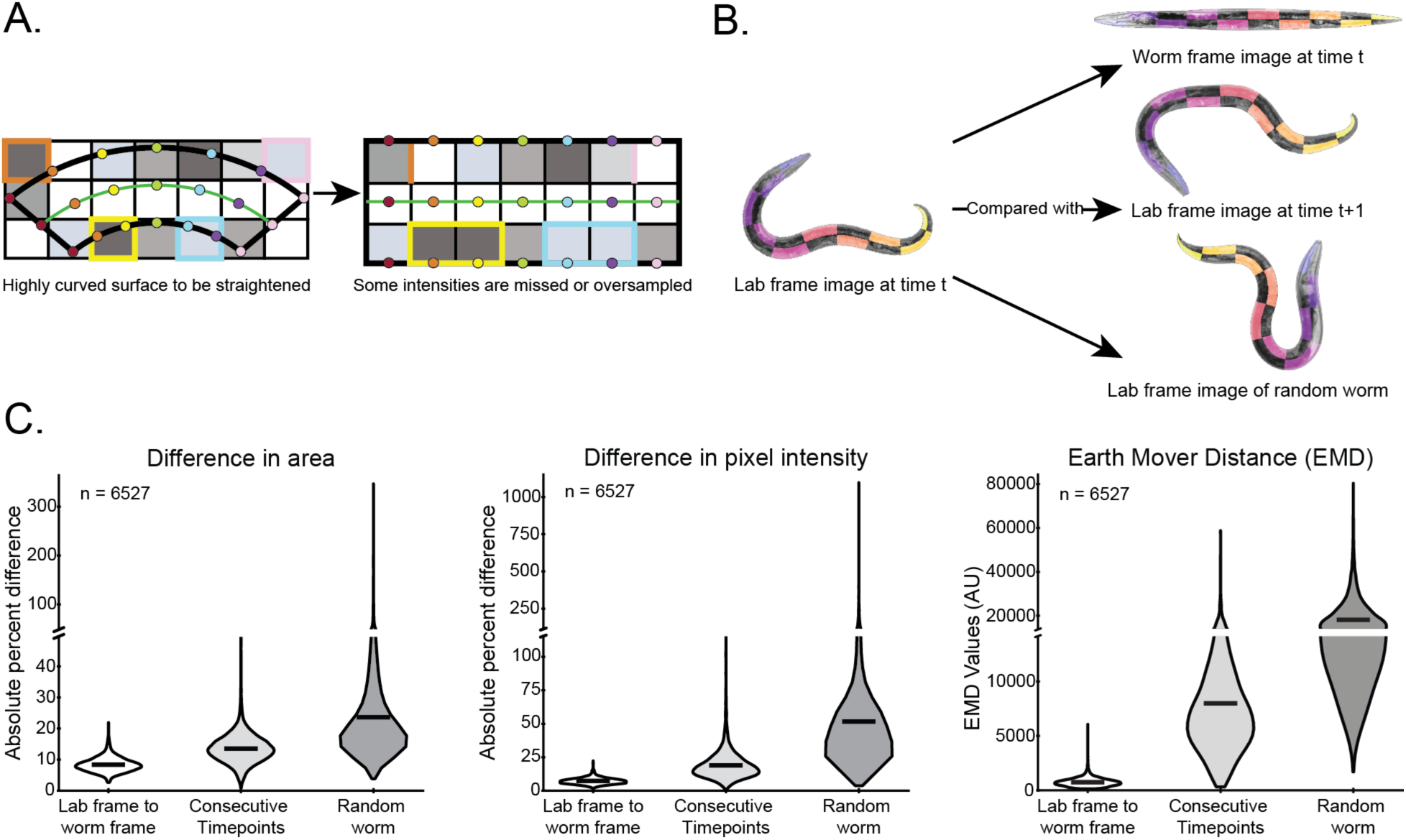
Quantifying the effects of straightening images into the worm frame of reference. (A) Illustration of image distortions that arise when straightening a curved shape. The curved shape on the left can be straightened by sampling across 7 evenly spaced points perpendicular to the centerline (green line), depicted by the colored dots. Points of the same color indicate the perpendiculars. To straighten, the intensity of the pixel each colored dot falls on in the original curved surface is mapped to the corresponding coordinate in the straightened image on the right, indicated by the corresponding colored dots. (In practice, image intensities for positions away from pixel centers are linearly interpolated across the neighboring pixels, reducing overt pixel-boundary effects. This does not eliminate the image distortions, however.) Across the top edge of the shape, which is longer than seven units in the left, “lab-frame” image, sampling at only seven positions leads to several pixels in the left image with intensities that are omitted in the ”worm-frame” image at right (undersampling). These pixels are outlined in orange and pink in the left image. Sampling the bottom edge, which is shorter than seven units in the left image, leads to two pixels being doubly-represented in the right-hand image (oversampling). These are outlined in yellow and blue. (B) Example of checkerboard sections used for analyzing the effects of over- and under-sampling. (Here each image is shown divided into 20 distinct sections for clarity; we used 40 sections for the full-resolution images we actually analyzed.) Pixel-wise differences are computed for matching sections and their absolute values summed. In this way, oversampling on the inside of curves and undersampling on the outside of curves cannot simply cancel out, allowing us to estimate the total absolute image distortion. (C) Quantification of the effects of computationally straightening 6527 *Plin4::GFP C. elegans* fluorescent images. For each of three difference measures, we examined the distribution of the absolute values of the differences, summed across checkerboard squares across three different image-to-image comparisons. At left, is the summed absolute differences in area of matching checkerboard regions: between the worm-frame and lab-frame images (left column); between lab-frame images at two consecutive timepoints (representing the degree of variability that might be expected between individuals in essentially identical biological states); and between two randomly-selected lab-frame images (representing the dissimilarity expected between entirely unrelated images). At center the summed absolute differences in total pixel intensity between matching checkerboard regions is shown. At right is a more sophisticated comparison of the differences between per-checkerboard-region pixel intensity histograms, the Earth Mover Distance. In all cases, the distortions induced by transforming an image from lab frame to worm frame are smaller than those produced from sequential images of the same individual in a different physical position.

Such issues are often waived away by noting that computationally straightening curved surfaces is an affine transformation, and is thus fully invertible (9, 16). And indeed, it is equally possible to map from the lab to worm frame and vice versa. This latter property is especially useful for making tissue-to-tissue correspondences between two different worm images. Since the origin of every pixel in the worm frame of reference is known, point-correspondences can be made between any lab-frame image and another image from the worm frame of reference. However, an invertible transformation does not mean that it is area-preserving, and as shown in Fig 2A. Particularly, the inner edges of curves are expanded during straightening and the outer edges are shrunken. This is of particular consequence for quantitative fluorescence imaging: there is no guarantee that total fluorescence intensity in highly curved sections of an animal will be conserved during the lab-frame to worm-frame transformation. We thus found it important to evaluate whether the size of this effect is a relevant practical problem.

We quantified the effects of computationally straightening *C. elegans* images using a dataset of several thousand (n=6527) fluorescence images of adult *Plin-4*::GFP*;spe-9(hc88)* animals (See Methods). We measured the differences in area, total pixel intensity, and the distribution of pixel intensities between fluorescence images before and after straightening (i.e. in the lab and worm frames of reference). In all cases these statistics were calculated only over pixels within the region corresponding to an individual *C. elegans*, ignoring background pixels. To understand the scale at which such differences would become biologically meaningful we compared these statistics to those calculated between unstraightened, lab-frame images of the same individuals taken three hours later. We reason that worm size and GFP expression does not vary greatly in adult *Plin4*::GFP animals over a three-hour interval. Thus, measured differences between such images would be due to variability in the imaging process or changes in the animal’s position/posture, and thus represent a level of variation that is safe to consider as “biologically negligible”.

Therefore, we can conclude that the errors produced by the straightening procedure are sufficiently small as to be biologically negligible if they are smaller than those observed between consecutive lab-frame images taken at three-hour intervals. To define a scale at which image differences are definitively non-negligible, we also compared lab-frame images of random pairs of different, non-age-matched individuals.

To measure the total amount of area and pixel intensity altered by the straightening procedure, it was important to ensure that worm area/pixel intensity created on the inside of each bend was not nullified by area/intensity removed on the outside of the same or other bends. Even if net image intensity or area remains unchanged, we wanted to examine the extent to which image measurements from any particular anatomical region might be distorted by straightening. Therefore, we split the worm regions into a “checkerboard” pattern to ensure that the inside and outside edges of each bend would be compared separately (Fig 2B). The size of the checkerboard was chosen to be smaller than the size of typical worm bends. The differences in area shown in Fig 2C are the sum of the absolute value of the difference in area of each matching checkerboard section between a worm-frame and lab-frame image, or two different lab-frame images. Similarly, the difference in total fluorescence intensity is the sum of the absolute differences in total intensity between matching sections.

Image regions with the same total fluorescence intensity do not necessarily have identical pixel patterns. For example, image warping might lead to the loss of a single very bright pixel and the duplication of several dimmer pixels, leading to an identical total pixel intensity between the warped and un-warped images. These images, however, would have very different distributions of pixel intensities. We therefore also compared how similar or dissimilar the distribution of intensities were across pairs of images, by first computing pixel intensity histograms for each image and then calculating the Earth Mover Distance (EMD, also known as Wasserstein Distance) metric to define the similarity of those histograms. This metric has been used to measure image histogram similarity for color-based image retrieval, texture metrics, and shape matching (27–33). The EMD calculates the amount of “transportation work” needed to transform one histogram into another, effectively penalizing differences in nearby histogram bins less than differences that require moving histogram “mass” between more distant bins (34, 35) (Fig S1). For this measure, the total reported EMD between a pair of worm images is the sum of each of the EMDs between matching checkerboard sections.

Using longitudinal brightfield and fluorescent image timeseries acquired using in our Worm Corral system, we selected images from 3–7 days post hatch (corresponding to young through mid-adulthood), resulting in a dataset of 6527 individual *C. elegans* images (36, 37) (See Methods). We defined three sets of image pairs to compare: lab-frame vs. worm-frame images; consecutive lab-frame images of the same individual; and random pairs of lab-frame images from different individuals. The worm-frame images were straightened as described above and the above measures of differences among image pairs were calculated. Overall the differences induced by the worm-straightening procedure were dramatically smaller than those between lab-frame images of the same individual taken three hours apart (Fig 2C; Two-tailed T-test of independent means p-values < 1e-30 in all cases). This finding suggests that the changes in image intensity induced by computationally straightening curved worm bodies are negligible compared to the other sources of biological and/or technical noise in the imaging system. This result confirms the viability and accuracy of performing quantitative analyses in the worm frame of reference.

### Accounting for internal variation

Although straightening the body accounts for differences in overall posture, *C. elegans* vary in size and shape, both in the same animal over time and among individuals. It is possible to standardize all worm-frame images to the same pixel length by adjusting the sample spacing in the AP dimension during the straightening procedure, uniformly compressing the worm- frame images of longer individuals and stretching those of shorter individuals. Similarly, the sample spacing along the DV axis can be adjusted to transform wider- or narrower-than-average individuals to a pre-specified width. Moreover, this DV sample spacing can be adjusted independently at each point along the AP axis to produce worm-frame images with a standardized nose to tail “width profile”. Lab-frame images from individuals of variable shapes and sizes can thus be standardized to enable direct comparison (Fig 3).

**Fig 3.**
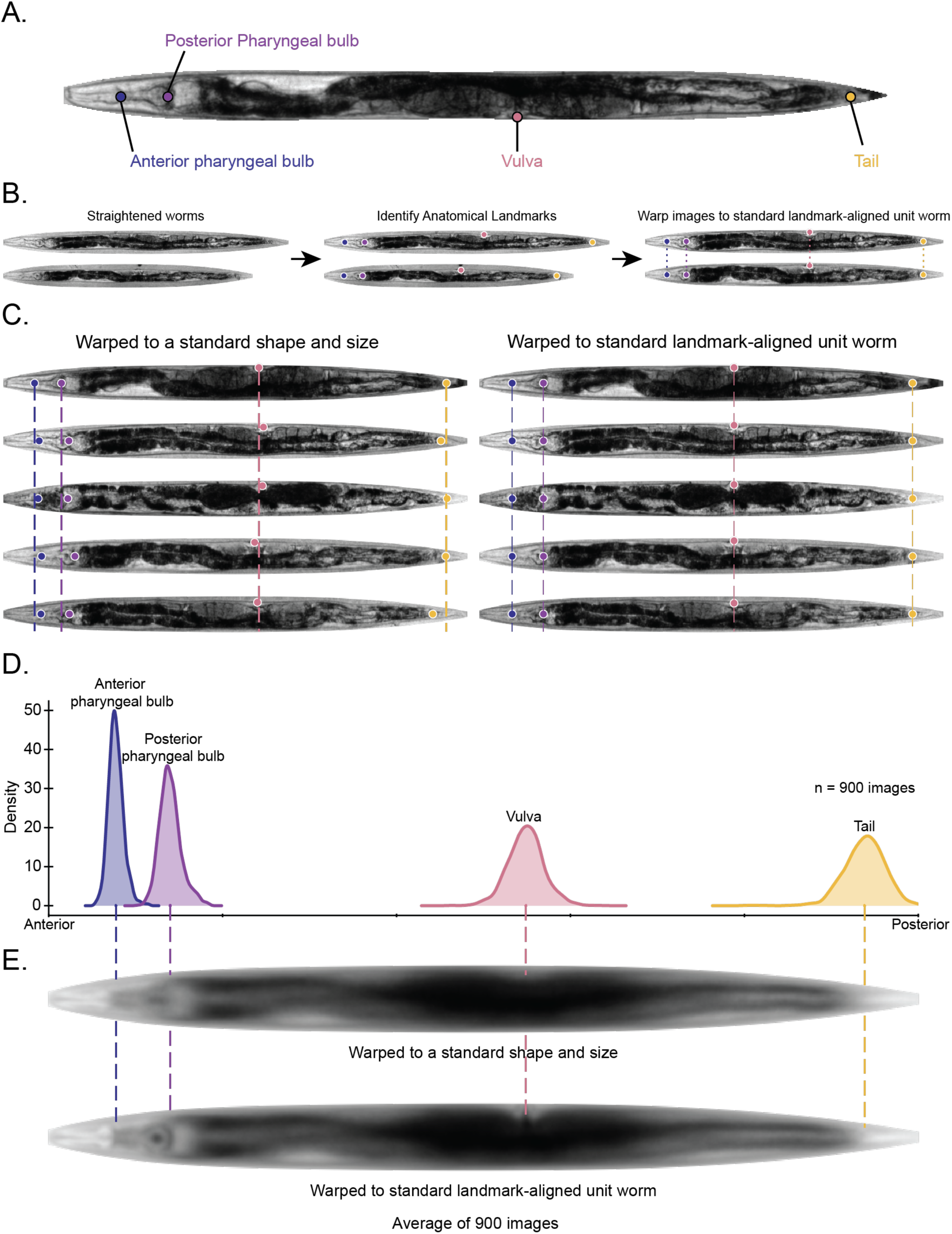
Correcting for individual variation with anatomical landmarks. (A) We identified four anatomical landmarks which can be reliably identified in whole-body brightfield images: the anterior and posterior pharyngeal bulbs, the vulva, and a “tail” landmark representing the posterior-most point with distinguishable internal structure. (B) These landmarks can be used to bring different worm-frame images into more precise register. First, anatomical landmarks are identified from worm-frame images. Next, the landmarks are brought into alignment by longitudinally warping the worm-frame images, to account for worm-to-worm internal variation. (C) Left: representative examples of worm-frame images of age-matched isogenic individuals warped to a standard size and shape with anatomical landmarks identified for each image. Dotted lines show how anatomical landmark locations can vary between individuals. Right: the same animals are shown after the landmarks are brought into alignment. (D) The distributions of the locations of each landmark across 900 brightfield images of isogenic adult animals are shown. Locations are depicted as the fraction along the anterior–posterior axis, rather than absolute pixel distances. (E) Top: the average of 900 brightfield worm-frame images, warped to a standard shape and size but without landmark alignment reveals only general anatomical structures, such as the pharynx and intestinal lumen. Bottom: the average of the corresponding landmark-aligned image shows considerably more anatomical detail.

However, shape standardization alone is not sufficient to account for individual variation because individuals differ in their precise relative location of internal structures as well. This is made apparent by highlighting the positions of a set of anatomical landmarks common to all wild-type adult *C. elegans*: the anterior and posterior pharyngeal bulbs, the vulva, and a “tail” landmark, which we define as the point after which there is no resolvable internal anatomical structure in our brightfield images (Fig 3A). This “tail” point, at which the worm becomes markedly more clear under brightfield imaging at the resolution employed in this study, can be reliably located despite not corresponding to a specific internal anatomical feature *per se*. Even in worm-frame images of age-matched isogenic individuals, warped to a standard size and shape, there continues to be variation in the location of those anatomical landmarks (Fig 3C, left). This indicates that, in addition to differences in posture, individual *C. elegans* are distinct in terms of the precise position of internal anatomical structures, which poses a further challenge in processing and analyzing such images.

To better account for inter-individual variation, we use the above anatomical landmarks as fiducial markers to bring images into closer alignment across individuals. Instead of evenly sampling the lab-frame image in the AP dimension, we calculate a variable sample spacing (see Methods) that smoothly warps worm-frame images such that the location of each landmark is placed at the population-average position (Fig 3B). For example, if an individual animal has a large distance between the posterior pharyngeal bulb and the vulva compared to the average worm, that portion of the lab-frame image will be sampled with a wider spacing to compress it into fewer pixels in the worm-frame image. Note that the sample spacing is computed using a smooth curve rather than linearly from landmark to landmark, to better model *C. elegans* as a compressible object and to avoid image discontinuities or artifacts at landmark positions (see Methods).

This longitudinal (AP dimension) warping ensures that every worm-frame image has approximately the same internal coordinates and allows coarse anatomical regions to be easily identified and compared across images. Manually annotating the anatomical landmarks of 900 brightfield adult *C. elegans* images allows us to quantify the distribution of landmark locations (Fig 3D). Without such landmarks, an average of these 900 brightfield images (warped to a standard length and width) reveals only general anatomical structures, such as the pharynx and intestinal lumen (Fig 3E, top). However, using the landmark positions to align the internal coordinates produces an average image with greater resolution of anatomical structures (Fig 3E, bottom). This illustrates how aligning images via anatomical landmarks can better account for internal variation and allow more precise comparisons among worm-frame images. Moreover, better internal coordinates can also improve the ability to make correspondences between different lab-frame images: for example, being able to define a region to compare across every lab-frame image as “the portion of each animal from the tip of its nose to the center of the anterior pharyngeal bulb” is more precise and biologically meaningful than comparing regions defined as “the portion of each animal from the tip of its nose and extending 150 microns posterior.”

### Automated identification of anatomical landmarks

Despite the utility of using anatomical landmarks as fiducial markers for image warping, manually entering the position of each landmark in each image (lab- or worm-frame) is infeasible for large image datasets. To address this, we developed a fully automated system to identify the location of these landmarks from worm-frame images, depicted in Fig 4A. We use a sequence of convolutional neural network models that take a brightfield worm-frame image as input and produce as output the AP coordinate of the four anatomical landmarks (Fig 4A). As *C. elegans* can crawl on either their left or right sides, we first apply a binary dorsal/ventral classifier to identify which side the animal is moving on (equivalently, whether the vulva appears on the top or bottom of the worm-frame image) and flip the image if necessary to ensure the same orientation for subsequent steps (38). We then employed a set of U-net convolutional neural networks (U-net CNNs) to transform brightfield images into a set of “keypoint map” images. These keypoint maps robustly encode the AP coordinate of each anatomical landmark in a way that is both easy to train CNNs to produce and easy to read out with simple processing (39). We examined two different forms of keypoint maps: a 1D gaussian “hot spot” placed over the AP position of the landmark, and a sigmoid function centered on the AP coordinate of the landmark (Figure 4B). Further details regarding the network and its training can be found in the Methods section.

**Fig 4.**
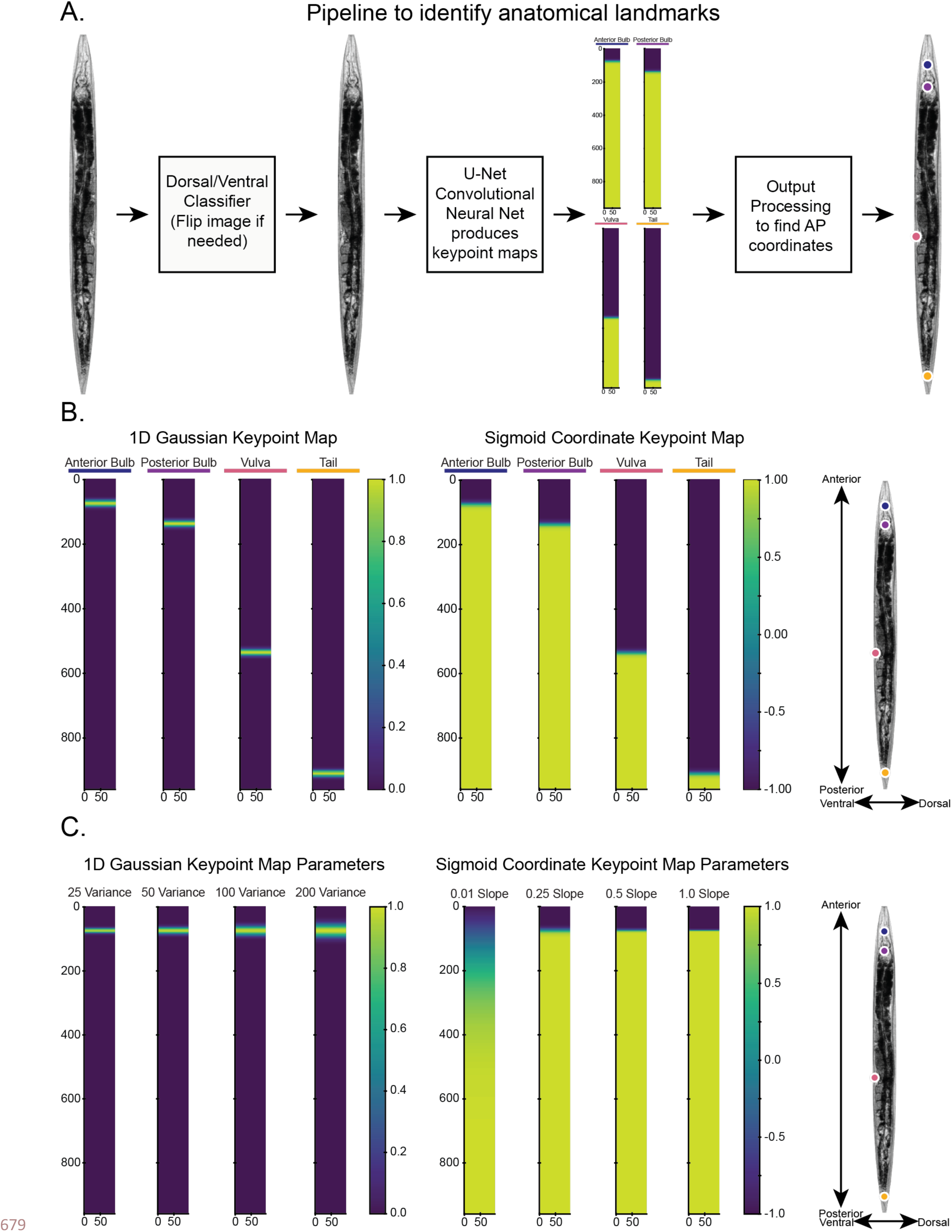
Automatic identification of anatomical landmarks. (A) Our overall process for identifying anatomical landmarks is as follows. Input lab-frame images are transformed into the worm frame of reference and warped to a standard shape and size (960x96 px). A Resnet-based convolutional neural network (CNN) classifier is used to determine the dorsal and ventral sides of the image, and images are flipped as needed to place the ventral side to the left. A U-net- based CNN is trained to produce output images that encode the location of each landmark (“keypoint maps”). These keypoint maps are then processed to recover the specific location of each landmark along the anterior–posterior (AP) axis. (B) We examined two schemes for encoding landmark locations in images, which the U-net classifier would be trained to reproduce. Left: Gaussian “hotspot” images in which the landmark location is the center of a 1- dimensional Gaussian function evaluated on the 2-dimensional image grid. Right: sigmoid images, in which the landmark location is the zero-crossing of a 1-dimensional sigmoid function (mapping from -1 to 1) evaluated on the 2-dimensional image grid. **(C)** We also examined the effects of varying the variance and slope parameters on the Gaussian and sigmoid images, respectively. Both parameters, illustrated here, control how “sharply” vs. “fuzzily” the keypoint map encodes the landmark location.

This approach to identifying anatomical locations via keypoint maps was influenced by the maps generated by DensePose, which establishes point correspondences between image pixels and a 3D model of the surface of the human body (40). Wang *et al.* also adopted this type of pixel-wise representation in their Celeganser segmentation tool, which encodes the AP and DV position of each pixel in a lab-frame image as the image intensity value at that position in a pair of output images (one for AP positions and one for DV positions) (12). Because identifying the location of a landmark only along the anterior-posterior-axis is sufficient for anatomical alignment between individuals, we trained our model only to identify the AP-coordinate of each landmark.

### Keypoint map representations can accurately locate anatomical landmarks

We evaluated the accuracy of our landmark-prediction CNNs by measuring the absolute distance (in pixels) between the predicted AP coordinate and the manually annotated ground-truth value, using a test set of images that were not used at any point in training the CNNs. We next compared the magnitude of these errors to the distances between the landmark positions in our ground-truth dataset and the positions of those same landmarks as annotated by other human raters (See Methods). In the best case, an automated system would predict landmark locations with no more imprecision than across human raters. In the worst case, a poorly trained model would not use the input image at all, and simply blindly guess that each landmark is at the average position across the training data.

We then examined how our automated system performed with respect to these best- and worst-case scenarios, across several different model parameters. For the gaussian keypoint maps, we examined different variance parameters, resulting in larger or smaller hotspots (Fig 4C). For sigmoid coordinates we varied the slope parameter, tailoring the sharpness of the transition between positive and negative values (Fig 4C).

We found that both 1D gaussian keypoint maps and sigmoid coordinates are robust, achieving accuracy as well as or better than human raters across a wide range of parameter values (Fig 5). As a difference of one pixel translates to 1.3 microns in our optical system, even the worst single error in the worst-performing model was less that 120 microns (or approximately one tenth of the length of an average adult animal) from its true position (Table S2). Based on these results we conclude that the sigmoid coordinate and 1D gaussian keypoint maps are sufficient representations of anatomical landmark location. The sigmoid coordinate map with a slope of 0.5 performed the best overall, with the lowest total error averaged across all landmarks (Table S2). We therefore used this scheme throughout the subsequent experiments and analyses.

**Fig 5.**
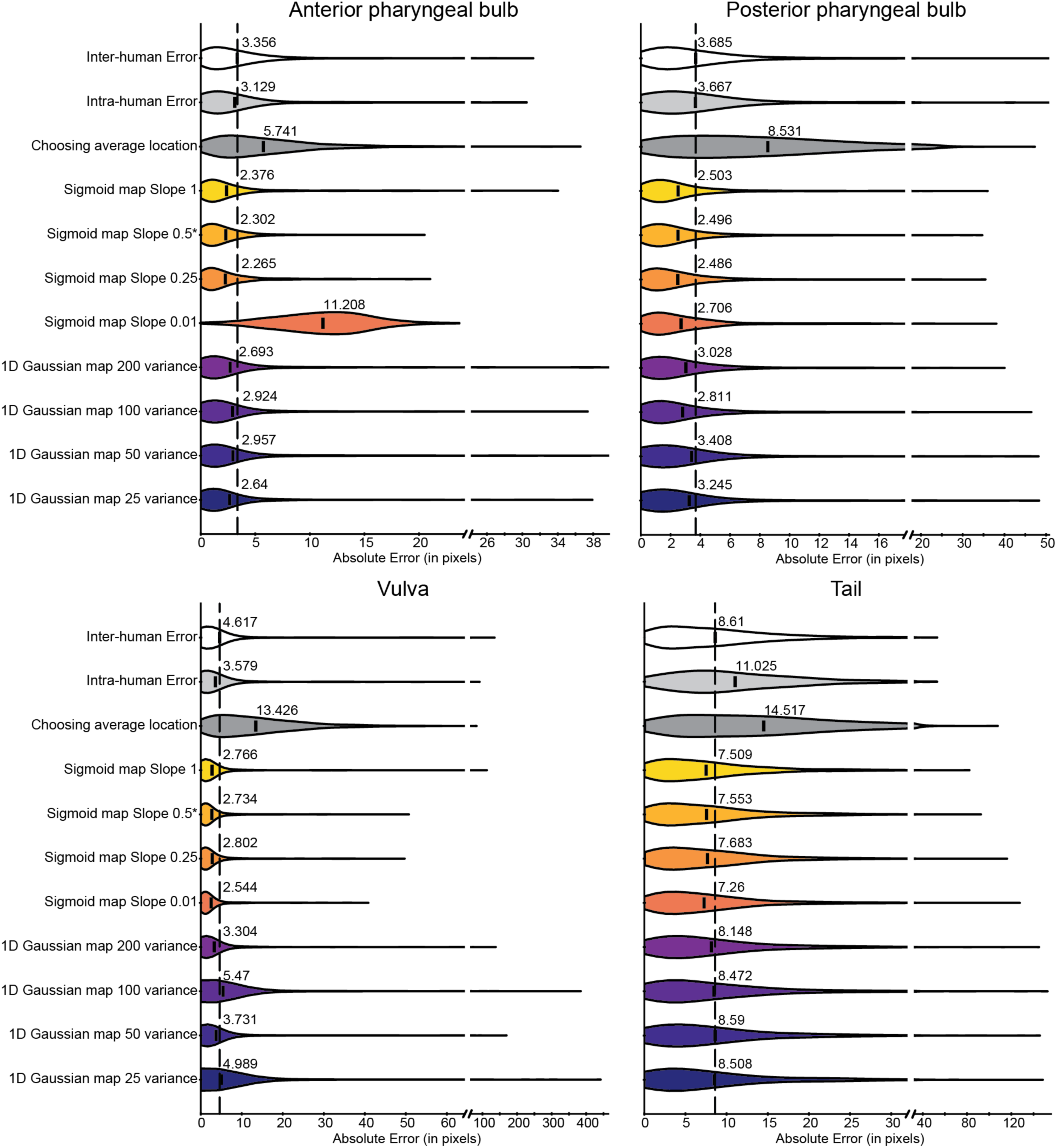
Keypoint map representations can accurately locate anatomical landmarks. Shown are the distributions of distances between “ground truth” landmark locations (produced by a single human rater) and the locations identified by other means. “Inter-human error” is the distribution of distances between the 200 ground truth locations and those provided by four other human raters. The mean value is marked with a dashed line for comparison with the other distributions. “Intra-human error” is the distribution of distances between the ground- truth locations and those provided by the same human rater several months afterward. “Choosing average location” is the distribution of errors produced by simply choosing the mean landmark location for each image, without reference to the image content at all. All other rows refer to errors produced by U-net CNNs trained to reproduce keypoint map images with the stated scheme (Gaussian or sigmoid) and parameter values. We found that the sigmoid coordinate keypoint map with a slope of 0.5 was the overall best, consistently outperforming even human raters on all landmarks.

Although the keypoint map representations were able to successfully be used to predict the four anatomical landmarks, some landmarks were easier for the model to identify than others. The two pharyngeal bulbs were the easiest anatomical landmarks for the model to locate while the vulva was the hardest. Visual inspection of the worst-localized landmark shows that the pipeline failed mostly on images where even human raters had difficulty identifying specific structures (Fig S2).

### Anatomical variation across individuals and over time

We next used these tools to examine how, at both the individual and population levels, *C. elegans* change morphologically with age. We straightened and identified the anatomical landmarks from brightfield images of isogenic, age-matched *Plin4::GFP;spe-9(hc88)* individuals between 2 and 6 days post-hatch (dph) (See Methods). Worm postures and anatomical landmark locations were manually corrected as needed. This dataset allowed us to examine early age-related phenotypes: at 2 dph, worms are young adults just beginning to lay eggs, while by 6 dph the shortest-lived individuals have died.

To visualize the variability in anatomical landmark locations across an age-matched population, we plotted a kernel density estimate of the distribution of landmark locations at 3.5 dph (Figure 6A, left). At this age, worms have reached their maximum length and remain generally healthy with high levels of mobility and physiological function (41–44). Moreover, at 3.5 dph, the population variation in landmark locations is the smallest compared to other ages (Fig S3, Table S3). All anatomical landmark distributions were approximately normal (Shapiro-Wilks p-values for normality all < 0.002). At this age, the tail position and worm length measurement were the most variable, and the anterior and posterior pharyngeal bulb positions the least variable. The anatomical landmark locations can also be presented normalized to the total worm length, effectively representing each locations as a of the distance along the AP axis (Fig 6B, right). The two pharyngeal bulbs are located at around 11% and 19% of the total length, respectively, while the average vulval position is at 50% of the worm length. The tail location (which as above does not refer to a specific anatomical structure but rather the last point of optical density visible at 10× magnification in our system) is also consistently around 94% of the worm length (Fig 6B right, Table S4). Using units of either absolute pixels or of relative percentages does not dramatically change the distributions (Fig 6A left vs. right); the coefficients of variation for each landmark distribution are very similar in either unit (Table S3 vs Table S4). This similarity results from both the low variability in length across the population and the overall stereotypy in relative landmark positions, and suggests that either set of units are appropriate for use.

**Fig 6.**
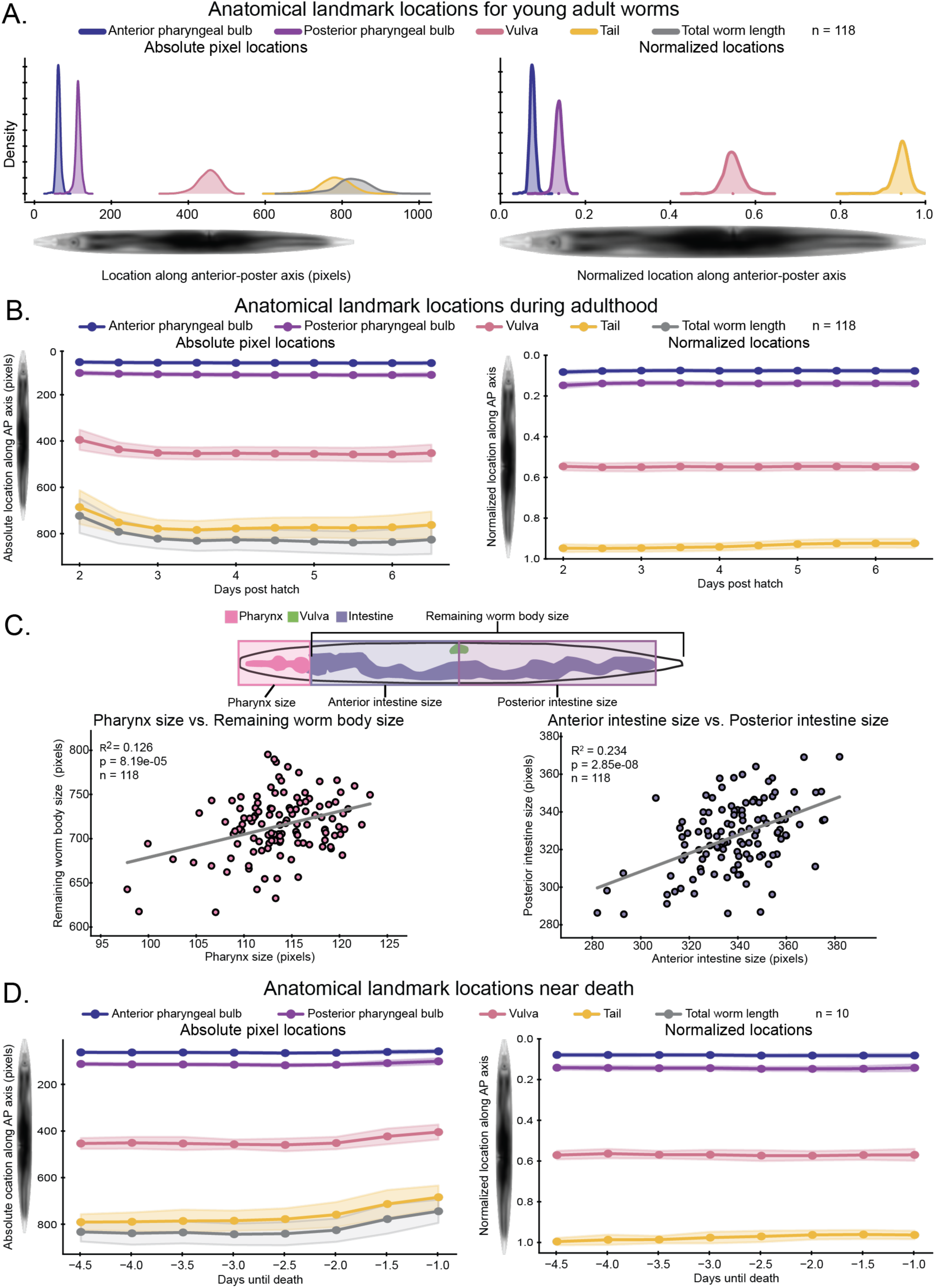
Individual and population anatomical variation. (A) Distributions of the anatomical landmark locations of isogenic age-matched *Plin4::GFP* animals at 3.5 days-post-hatch, in absolute (left) and relative (right) units. (B) Population averages of anatomical landmark locations 2–6.5 days-post-hatch. Standard deviations are shown in shaded regions. Left: absolute pixel locations, which show a previously observed early-adulthood growth phase between 2-4 days-post-hatch. Right: relative landmark locations (normalized by total worm length) show that there is no change in anatomical landmark positions relative to one another and to the animals’ length, suggesting that early-adult growth is proportional across the whole body. (C) Left: pharynx size is not strongly correlated to the remaining body size in animals 3.5 days-post-hatch, suggesting that the size of the pharyngeal and gonad/intestinal body compartments are not tightly coupled. Right: anterior and posterior gonad/intestine (defined by the location of the vulva) are also not strongly correlated, suggesting that there is not tight control over the relative position of the vulva within that compartment. (D) Population averages of anatomical landmark locations for short-lived individuals from 4.5 days to 1 day prior to death. Left: absolute pixel locations show a characteristic pre-death decrease in worm length Right: normalizing by total worm length shows there is no change in relative landmark locations despite this shrinkage, that it is due to a proportional decrease in size across all body compartments.

We next examined how anatomical landmark locations change over time. *C. elegans* increase in size through early adulthood (41–43) (Figure 6B left). During this young adult growth, the average anatomical landmark positions track with total length. Overall, the relative anatomical positions remain constant when normalized to total worm length, despite the increase in total worm size throughout young adulthood (Figure 6B right). This consistency suggests that all the anatomical compartments of *C. elegans* that we measured grow proportionally in the first few days of adulthood. If this were not the case, and, for example, the intestine was the site of the most growth, then one would expect that the *relative* position of the anterior and posterior pharynges would shrink proportionally. The flatness of lines in Figure 6B (right) indicates that, instead, growth is evenly distributed across the entire animal.

We next asked whether deviations from the average size of each anatomical compartment are correlated across individuals. This would provide evidence as to whether the developmental programs that determine each compartment’s size are coupled vs. proceed largely independent of one another. If, for example, the pharynx and intestine/gonad sizes were determined by coupled processes, then larger-than-average worms would typically have both pharynges and gonads/intestines that were larger than average. Alternatively, if the sizes of these compartments were determined by independent processes, above-average body length could be driven by above-average pharyngeal size, above-average intestinal and/or gonad size, or both (at a correspondingly lower rate). To address this question, we examined whether pharyngeal size was correlated with the size of the rest of the body, mostly determined by the intestinal and gonad size (Fig 6C, top). The correlation between pharyngeal and remaining body length was modest at best (Fig 6C, left), suggesting that there is only a weak coupling between the developmental programs that determine pharynx size and those that determine the sizes of the intestine, gonad, and other organs.

We also examined the related question of where the vulva is placed within the intestine/gonad. The intestine/gonad compartment is defined as the region between the posterior pharyngeal bulb and the tail landmarks (Fig 6C, top). We examined vulval placement within this region by comparing the lengths of the intestine/gonad anterior to and posterior to the vulva. If the placement of the vulva was tightly coupled with the rest of gonad and intestinal development, we might expect to see that the length of that compartment anterior and posterior to the vulva would vary in fixed proportion. We also only observed a modest correlation between these lengths (Fig 6C, right). Overall, these results suggest that there is neither tight developmental coupling between the relative scaling of the pharynx compared to the remaining body (Fig 6C, left) nor the position of the vulva with respect to the intestinal/gonad compartment (Fig 6C, right).

A subset of individuals died or came to within 1 day of death during the timeline of our dataset (2–6 days post-hatch). Near-death changes in body volume have been observed previously, with a sharp decrease in body size about a day prior to death followed by a recovery of pre-death size after death (3, 44). We observed this decrease in body length among the short-lived subset of individuals, beginning approximately two days prior to death (Fig 6D, left). We next examined whether this shrinkage occurs only in one or another set of tissues and/or organs. The largest shrinkage in absolute terms occurs in the intestinal/gonad compartment (Fig 6D, left); however, that also represents the largest compartment in the worm body. By expressing anatomical landmarks in terms of their relative position from anterior to posterior (Fig 6D, right), it is clearly visible that there is effectively no change in the relative position of each anatomical landmark during near-death shrinkage. The decrease in size is proportional across the entire body, with each region between pairs of landmarks shrinking by almost the same degree as each other.

## Discussion

With many high-throughput *C. elegans* culturing systems now available, a key challenge is analysis and automated interpretation of the large image datasets that such systems can produce. One major barrier to automated comparisons among many individual *C. elegans* is their flexible and deformable nature. This positional and anatomical variation among individuals must be accounted for to compare brightfield or fluorescent image patterns across the population and/or over time. Here we describe an automated pipeline to perform these tasks in whole-body images of adult *C. elegans,* without the need for fluorescent fiducial markers (from cellular stains or transgenic reporter animals). This machine-learning approach can be applied to fluorescence or other imaging modalities provided there is a corresponding brightfield image. Further, we propose that the underlying neural network could also be retrained to identify landmarks directly from many different classes of fluorescence images.

An important step in developing machine-learning driven image analysis tools is to determine the best data representation on which to train. We found empirically that regression techniques to transform an input image directly into a list of (x, y) coordinates for each image landmark, which have been successfully applied on images of human faces and other similar tasks (45, 46), did not function well in this setting. Instead, we found that image-to-image regression where the output was not the coordinates of the location of the landmark, but instead an image that implicitly and robustly encoded that location, was more successful. Ultimately, this approach was able to predict anatomical landmarks with the same accuracy as experienced human annotators, but with substantially reduced time and effort.

In this work, we demonstrated how automatically identified landmarks can be used for basic morphometric tasks, examining growth, shrinkage, and organ/tissue scaling. However, these tools have wider applicability. Specifically, accounting for internal anatomic variability can greatly improve the fidelity of fluorescent image analyses. As depicted in Figure 3, removing both positional and anatomical variation allows for greater resolution of anatomical structures when averaging images. The same would be true for fluorescent images, both in terms of calculating average images and performing principal components on images to calculate correlated modes of variation in fluorescent intensity around those averages.

In addition, an average brightfield worm image can be used to manually identify specific regions or tissues of interest, creating an atlas of regions which can be used for tissue-specific analyses (Fig S4). Specifically, labeled regions from the atlas can be propagated to accompanying fluorescent images, either in the worm frame of reference, or mapped back into the lab frame of reference to analyze tissue-specific expression. Finally, after removing positional and internal anatomical variation, images in the worm frame of reference can be used to examine morphological changes between individuals and across time from brightfield images or to investigate spatiotemporal gene expression patterns from fluorescent images. Overall, correcting for positional and anatomical variation, as described in this work, can improve the fidelity of many different classes of *C. elegans* imaging tasks.

## Methods

### *C. elegans* strains and single-animal culturing system

We obtained VT1072 (*Plin-4*::GFP) animals from the Caenorhabditis Genetics Center (CGC), which is funded by NIH Office of Research Infrastructure Programs (P40 OD010440), and crossed them into the temperature-sensitive sterile mutant BA671 (spe-9(hc88)) (47). Longitudinal brightfield timeseries images of *C. elegans* animals were acquired using the single-animal culturing system described by Pittman et al (36). Brightfield and fluorescent images were acquired in succession every 3-6 hours from hatching until death. Timepoint images were taken from multiple experiments where animals were cultured at 20°C or 25°C in the worm corral. At 20°C, reproduction was prevented through the use of *pos-1* RNAi (36, 48). For worms cultured at 25°C, the *spe-9* mutation prevents fertilization. All anatomical variation experiments were performed using animals cultured at 25°C.

### Worm finding and straightening

To identify centerlines from brightfield images, we applied a deep-learning based image segmentation tool (12) to find both the set of pixels corresponding to a single individual, termed the “pixel mask”, and estimates of the anterior-posterior and medio-lateral coordinates for each pixel within this mask. To smooth these estimates, nose-to-tail centerlines of the animals were identified from the mask using a thinning process called skeletonization, and separately from the pixel-wise coordinate maps by tracing the local minima of the medio-lateral coordinates, corresponding to the centerline. The best option was chosen and manually modified, as necessary, by a human operator. The centerline was then fit to a 3^rd^ degree parametric spline x(c), y(c) representing the (x, y) coordinate in the original image of the point c pixels along the centerline from nose to tail. The tangent to the spline was calculated via first derivatives and then rotated 90° to define the local direction of the dorsal–ventral axis (though which side is dorsal vs. ventral is determined later; see below). Specifically, this normal vector at point c is defined as (-yʹ(c), xʹ(c)). At each pixel along the midline, the distance from the midline to the nearest edge of the pixel mask was calculated via the Euclidian distance transform, defining a “width profile” of each individual w(c) that provides the width in pixels of a given individual at point c. These profiles were smoothed by fitting to a 3^rd^ degree spline.

To straighten an image from the original “lab frame of reference” into the “worm frame of reference,” we calculated the centerline coordinates (x(c), y(c)) and normal vector (-yʹ(c), xʹ(c)) at one-pixel intervals from nose to tail, defining the length of the worm-frame image. For a 2k+1 pixel-wide image, for each point c we calculated the image coordinates along single-pixel steps from -k to k pixels out from the centerline at point c, in the direction of the normal vector. Thus, at each point c along the anterior-posterior axis of the worm we calculated the (x, y) coordinates in the lab-frame image of a strip of 2k+1 pixels running in the local dorsal-ventral direction. The lab-frame image is then sampled at each of these coordinate positions, producing a single column of pixels in the worm-frame image.

To produce worm-frame images with a standardized length and width profile that matches that of the “average” worm, we sampled the lab-frame image with a different number of steps along the centerline to “stretch” or “shrink” a given individual to a longer or shorter length, and/or along the normal at any given point to stretch or shrink the width of the worm at that point to match the average width profile. To warp the internal coordinates of an individual such that a particular anatomical landmark at a position of c1 pixels along the centerline instead falls at a new position c2 along the centerline, we fitted pairs of (*coordinate-in*, *coordinate-out*) points to a Piecewise Cubic Hermite Interpolating Polynomial (49), as implemented in the SciPy software package, using the points (0, 0), (*c1, c2*), (*length, length*) as input (50). We then calculated the *coordinate-out* values for evenly spaced *coordinate-in* values from 0 to *length*, and evaluated both the centerline spline *(x(c), y(c))* and normal spline along the now-unevenly-spaced *coordinate-out* values. Multiple landmarks required more pairs of (*coordinate-in*, *coordinate-out*) points to be fit to the polynomial.

### Landmark data collection and validation

To train and validate the CNN models, we selected images corresponding to adult timepoints from the image dataset described above, resulting in a dataset of 6527 total images from 155 individual *C. elegans* animals aged between 2 and 32.5 dph (or until death).

Centerlines and widths were identified from each brightfield image and straightened using the method described above for inages between 2 and 6 dph. The (*x,y*) coordinates of four anatomical landmarks (anterior and posterior pharyngeal bulbs, vulva, and tail) were manually labeled by an experienced *C. elegans* researcher using custom annotation software. These annotations were used as the “ground-truth” in training the CNNs. Ground truth dorsal/ventral classes for each worm image were established by the y-coordinate of the vulva, i.e. whether it was above or below the centerline.

The criteria for annotating the anatomical landmarks were as follows:

- Anterior pharyngeal bulb: the point on the animal’s midline at the most posterior end of the anterior pharyngeal bulb was selected.
- Posterior pharyngeal bulb: the midline point at the posterior end of the posterior pharyngeal bulb.
- Vulva: the point on the animal’s side at the middle of the vulva.
- Tail: the end of visible tissue/texture where the worm tissue goes from more textured to clear in our image data at the posterior end of the worm, generally slightly past the anus, which is not always directly visible in our images.

We randomly grouped these annotated images into “train”, “validation”, and “test” datasets (4515 “train” images, 1322 “validation” images, and 690 “test” images). These datasets were used to train, evaluate, and benchmark the neural networks, respectively. During the training process, the validation dataset was used to ensure the neural networks were not overfitting to the training data. The test dataset, however, was never used during the training process and was only used to benchmark the neural network performance. We ensured that all images from the same individual were grouped into the same datasets, to prevent overfitting to the idiosyncrasies of particular individuals.

### Checkerboard mask generation and analyses

“Checkerboard” mask regions were generated using full-body worm masks in the worm frame of reference. The worm masks are sliced into 10 equal segments along the length of the worm. Each region is further split along the centerline to create one dorsal and one ventral region, for 20 total. The checkerboard regions are then transformed back into the lab frame of reference to compare regions in the lab frame vs. worm frame.

Corresponding regions in the lab frame vs. worm frame were compared by calculating the difference in area, the difference in mean pixel intensity, and the Earth Mover Distance (Wasserstein distance) between pixel intensity histograms. To aggregate across all regions for a given image, we computed the sum of the absolute values of each of these differences/distances. All measurements were performed on flatfield-corrected images and were masked so that only worm-associated pixels went into the analyses.

### Network architecture and training

For the Dorsal/Ventral classifier we used ResNet34 with a binary output layer. We used a cross entropy loss function when training as implemented by PyTorch (nn.CrossEntropyLoss). We trained the classifier for 25 epochs with a batch size of 5 and saved the model with the lowest error on the “validation” dataset at the end of the training epochs. We used the Adam optimizer with a base learning rate of 0.0005, decreasing by half every 4 epochs. We performed this training scheme eight times and chose the model that stochastically had the lowest error for the final pipeline. The final Dorsal/Ventral classifier achieved an average accuracy of 96.12% on the “test” images (data not shown). Note that the model was neither trained on any images from the “test” dataset, nor were those images used to select among different models.

For the anatomical landmark prediction models we adapted the “Coordinate Regression” neural network architecture described by Wang et al. (12). The model has a U-Net architecture with each step in the U implemented as a ResNet34 encoder and a decoder with skip connections to the encoder and upsampling layers, resulting in a single output image that is the same size as the input (12, 39). To train the model we used a per-pixel L1 loss, calculated only on pixels corresponding to the “worm” image regions. Like Wang et al., we calculated the loss at each skip connection layer with multiple scales, again masked to include only worm image pixels (12). For each image we summed the loss calculated over all scales. We trained independent models for each landmark, resulting in four landmark models for the full pipeline. Each anatomical landmark prediction model was trained for 25, 50, and 150 epochs (data not shown for 50 and 150 epochs) with the same batch size and base learning rate as above. Unlike the Dorsal/Ventral classifier, we only trained the models a single time per parameter change. For the 50 and 150 epoch training schemes the learning rate decreased by half every 5 and 15 epochs respectively. All input and subsequently output images were 960 x 96 pixels.

All CNNs were implemented in PyTorch and trained on a single NVIDIA 1080TI.

### Keypoint map generation

“Keypoint map” images served as the target images that the U-Net was trained to produce based on input brightfield images. The original anatomical landmarks were specified by a human annotator on a worm-frame-of-reference image, in terms of pixels along the anterior– posterior (AP) axis, and pixels dorsal–ventral (DV) to the midline. The AP coordinate of each landmark was normalized to our standard worm length (960 pixels) by multiplying by a factor of 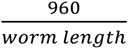, where the length of each individual worm was calculated as the total arc length of the centerline spline. For the vulva keypoint map, the DV coordinate of the landmark was set to the width of our standard worm at the calculated AP coordinate. For the other landmarks, the DV coordinate was set to zero (i.e. the centerline). Given the AP and DV coordinates for each landmark, the keypoint map images were generated as follows.

#### Gaussian keypoint maps

We used the SciPy stats package to generate a normal distribution centered at the anatomical locations in the keypoint map coordinate system with the variances specified in the text.

#### Sigmoid keypoint maps

To get the sigmoid centered at the anatomical landmark AP coordinate, we first generated a grid of the indices with the shape of the input image and then subtracted the landmark AP coordinate from this grid. This ensured that the landmark location would be at zero. Then we applied a sigmoid function to the index grid such that the pixel intensity value *I*_x_ of the keypoint map at pixel (x,y) in the standard worm coordinates is given by:

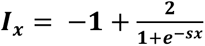

where s is the slope parameter specified in the text.

### Anatomical location keypoint map processing

We first smoothed the output keypoint maps produced by the U-Net by applying a gaussian filter with a standard deviation of one pixel and removed negative values by setting any pixels less than half of the smoothed maximum value to zero. For the gaussian keypoint maps, we employed the simple moment method described by Anthony and Granick (51) to identify the center-point of the gaussian in a fashion more robust than simply selecting the location of the maximum pixel. With this method, the AP coordinates of the output landmark position, *xc,* are calculated as:

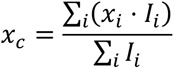

where *x_i_* is the AP coordinate of pixel *i* and *I_x_* is the intensity of the pixel at a given index (51).

For the sigmoid coordinate output maps, we first transformed the pixel intensities into a range suitable to calculate landmark positions using the moment method. Specifically, we transformed image intensities to place the landmark, originally where the sigmoid map is zero-valued, at the maximum intensity instead: *I_out_* = *M* − |*I*_in_|, where M is the maximum pixel intensity across the image. Essentially, this changes the sigmoid coordinate output map into a hotspot map. After this transformation, we identified the AP coordinate as described above.

To convert the AP coordinates from the standard worm (960 pixels long, with a standardized width profile) to those of any specific individual, we multiply by 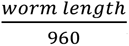, where *worm length* is calculated as above.

### Inter-rater agreement

As a benchmark, we compared the distance between the human-annotated ground-truth landmark locations and either the locations generated by our neural network models or those annotated by other human raters. We asked 3 raters well versed in *C. elegans* anatomy to identify the four anatomical landmarks in 200 worm-frame images sampled from the full dataset. As worms age their organs become more visibly disordered, making it increasingly difficult to identify the anatomical landmarks. The 200 images were thus equally sampled from 10 age bins ranging from young to old adulthood to ensure images from all ages were represented.

To account for systemic bias in each human rater’s annotations (e.g. perhaps one rater systematically annotates the pharynges as ten pixels anterior of where another would), we first calculated a bias term, defined as the average signed distance between a rater’s annotations and the ground-truth landmark coordinates for each anatomical landmark. This bias term was subtracted from the human rater’s annotated anatomical landmark locations before calculating absolute error metrics. The absolute error was then calculated as the distance between the AP- coordinates of the bias-corrected human-annotated landmarks and the ground-truth landmarks. This absolute error was averaged across all raters to obtain the “human error” shown in the figures.

## Code Availability

Source code is available as a part of the open-source Python package Elegant: https://github.com/zplab/elegant. Pose estimation code and trained model parameters are

part of the “Cegmenter” module. Anatomical landmark identification code and trained models are included in the “Keypoint Annotation” module.

## Acknowledgements

We thank Holly Kinser and Matthew Mosley for use of their worm corral images. We would also like to thank Nick Jensen and Matthew Mosley for their expertise in worm anatomy and help in generating the human rater benchmarks. Author N. Laird would like to thank the Meow Brothers Paulo and João for their unconditional love and snuggles. This work was supported by NIH grants T32 HG000045 and R01 AG057748, and a Beckman Young Investigator award from the Arnold and Mabel Beckman Foundation. Some strains were provided by the CGC, which is funded by the NIH Office of Research Infrastructure Programs (P40 OD010440).

## References

1. Churgin MA, Jung S-K, Yu C-C, Chen X, Raizen DM, Fang-Yen C. Longitudinal imaging of Caenorhabditis elegans in a microfabricated device reveals variation in behavioral decline during aging. Sengupta P, editor. eLife. 2017 May 24;6:e26652.

2. Zhang WB, Sinha DB, Pittman WE, Hvatum E, Stroustrup N, Pincus Z. Extended twilight among isogenic C. elegans causes a disproportionate scaling between lifespan and health. Cell Syst. 2016 Oct 26;3(4):333–345.e4.

3. Stroustrup N, Ulmschneider BE, Nash ZM, López-Moyado IF, Apfeld J, Fontana W. The Caenorhabditis elegans Lifespan Machine. Nat Methods. 2013 Jul;10(7):665–70.

4. Chung K, Zhan M, Srinivasan J, Sternberg PW, Gong E, Schroeder FC, et al. Microfluidic chamber arrays for whole-organism behavior-based chemical screening. Lab Chip. 2011 Nov 7;11(21):3689–97.

5. Kopito RB, Levine E. Durable spatiotemporal surveillance of Caenorhabditis elegans response to environmental cues. Lab Chip. 2014 Jan 21;14(4):764–70.

6. Keil W, Kutscher LM, Shaham S, Siggia ED. Long-term high-resolution imaging of developing C. elegans larvae with microfluidics. Dev Cell. 2017 Jan 23;40(2):202–14.

7. Hulme SE, Shevkoplyas SS, McGuigan AP, Apfeld J, Fontana W, Whitesides GM. Lifespan-on- a-chip: microfluidic chambers for performing lifelong observation of C. elegans. Lab Chip. 2010 Mar 7;10(5):589–97.

8. Aerni SJ, Liu X, Do CB, Gross SS, Nguyen A, Guo SD, et al. Automated cellular annotation for high-resolution images of adult Caenorhabditis elegans. Bioinformatics. 2013 Jul 1;29(13):i18–26.

9. Long F, Peng H, Liu X, Kim SK, Myers E. A 3D Digital Atlas of C. elegans and Its Application To Single-Cell Analyses. Nat Methods. 2009 Sep;6(9):667–72.

10. Nagy S, Goessling M, Amit Y, Biron D. A Generative Statistical Algorithm for Automatic Detection of Complex Postures. PLoS Comput Biol. 2015 Oct 6;11(10):e1004517.

11. Moore BT, Jordan JM, Baugh LR. WormSizer: High-throughput Analysis of Nematode Size and Shape. PLOS ONE. 2013 Feb 22;8(2):e57142.

12. Wang L, Kong S, Pincus Z, Fowlkes C. Celeganser: Automated Analysis of Nematode Morphology and Age. In 2020 [cited 2021 Jan 6]. p. 968–9. Available from: https://openaccess.thecvf.com/content_CVPRW_2020/html/w57/Wang_Celeganser_Automated_Analysis_of_Nematode_Morphology_and_Age_CVPRW_2020_paper.html

13. Wählby C, Riklin-Raviv T, Ljosa V, Conery AL, Golland P, Ausubel FM, et al. RESOLVING CLUSTERED WORMS VIA PROBABILISTIC SHAPE MODELS. Proc IEEE Int Symp Biomed Imaging Nano Macro IEEE Int Symp Biomed Imaging. 2010 Jun 21;2010(14-17 April 2010):552–5.

14. Raviv TR, Ljosa V, Conery AL, Ausubel FM, Carpenter AE, Golland P, et al. Morphology- Guided Graph Search for Untangling Objects: C. elegans Analysis. Med Image Comput Comput-Assist Interv MICCAI Int Conf Med Image Comput Comput-Assist Interv. 2010;13(Pt 3):634–41.

15. Hebert L, Ahamed T, Costa AC, O’Shaughnessy L, Stephens GJ. WormPose: Image synthesis and convolutional networks for pose estimation in C. elegans. PLoS Comput Biol. 2021 Apr 27;17(4):e1008914.

16. Peng H, Long F, Liu X, Kim SK, Myers EW. Straightening Caenorhabditis elegans images. Bioinformatics. 2008 Jan 15;24(2):234–42.

17. Christensen RP, Bokinsky A, Santella A, Wu Y, Marquina-Solis J, Guo M, et al. Untwisting the Caenorhabditis elegans embryo. eLife. 4:e10070.

18. Bao Z, Murray JI, Boyle T, Ooi SL, Sandel MJ, Waterston RH. Automated cell lineage tracing in Caenorhabditis elegans. Proc Natl Acad Sci U S A. 2006 Feb 21;103(8):2707–12.

19. Bubnis G, Ban S, DiFranco MD, Kato S. A probabilistic atlas for cell identification. ArXiv190309227 Q-Bio [Internet]. 2019 Mar 21 [cited 2022 Jan 26]; Available from: http://arxiv.org/abs/1903.09227

20. Chaudhary S, Lee SA, Li Y, Patel DS, Lu H. Graphical-model framework for automated annotation of cell identities in dense cellular images. eLife. 10:e60321.

21. Yemini E, Lin A, Nejatbakhsh A, Varol E, Sun R, Mena GE, et al. NeuroPAL: A Multicolor Atlas for Whole-Brain Neuronal Identification in C. elegans. Cell. 2021 Jan;184(1):272–288.e11.

22. Varol E, Nejatbakhsh A, Sun R, Mena G, Yemini E, Hobert O, et al. Statistical Atlas of C. elegans Neurons. In: Martel AL, Abolmaesumi P, Stoyanov D, Mateus D, Zuluaga MA, Zhou SK, et al., editors. Medical Image Computing and Computer Assisted Intervention – MICCAI 2020. Cham: Springer International Publishing; 2020. p. 119–29. (Lecture Notes in Computer Science).

23. Nejatbakhsh A, Varol E, Yemini E, Venkatachalam V, Lin A, Samuel ADT, et al. Extracting neural signals from semi-immobilized animals with deformable non-negative matrix factorization [Internet]. Neuroscience; 2020 Jul [cited 2022 Jan 26]. Available from: http://biorxiv.org/lookup/doi/10.1101/2020.07.07.192120

24. Iglesias JE, Sabuncu MR. Multi-Atlas Segmentation of Biomedical Images: A Survey. Med Image Anal. 2015 Aug;24(1):205–19.

25. Oliveira FPM, Tavares JMRS. Medical image registration: a review. Comput Methods Biomech Biomed Engin. 2014;17(2):73–93.

26. Qu L, Long F, Liu X, Kim S, Myers E, Peng H. Simultaneous recognition and segmentation of cells: application in C.elegans. Bioinformatics. 2011 Oct 15;27(20):2895–902.

27. Nayyeri F, Nasrudin MF. Image Matching Using Dimensionally Reduced Embedded Earth Mover’s Distance. J Appl Math. 2013 Dec 4;2013:e749429.

28. Peleg S, Werman M, Rom H. A unified approach to the change of resolution: space and gray-level. IEEE Trans Pattern Anal Mach Intell. 1989 Jul;11(7):739–42.

29. Rubner Y, Tomasi C. Texture metrics. In: SMC’98 Conference Proceedings 1998 IEEE International Conference on Systems, Man, and Cybernetics (Cat No98CH36218). 1998. p. 4601–7 vol.5.

30. Snow M, Van lent J. Monge’s Optimal Transport Distance for Image Classification. ArXiv161200181 Cs Math [Internet]. 2018 Apr 8 [cited 2022 Jan 28]; Available from: http://arxiv.org/abs/1612.00181

31. Ling H, Okada K. An Efficient Earth Mover’s Distance Algorithm for Robust Histogram Comparison. IEEE Trans Pattern Anal Mach Intell. 2007 May;29(5):840–53.

32. Cohen S, Guibasm L. The Earth Mover’s Distance under transformation sets. In: Proceedings of the Seventh IEEE International Conference on Computer Vision. 1999. p. 1076–83 vol.2.

33. Grauman K, Darrell T. Fast contour matching using approximate earth mover’s distance. In: Proceedings of the 2004 IEEE Computer Society Conference on Computer Vision and Pattern Recognition, 2004 CVPR 2004. 2004. p. I–I.

34. Rubner Y, Tomasi C, Guibas LJ. A metric for distributions with applications to image databases. In: Sixth International Conference on Computer Vision (IEEE Cat No98CH36271). 1998. p. 59–66.

35. Werman M, Peleg S, Rosenfeld A. A distance metric for multidimensional histograms. Comput Vis Graph Image Process. 1985 Dec 1;32(3):328–36.

36. Pittman WE, Sinha DB, Kinser HE, Patil NS, Terry ES, Plutzer IB, et al. A Simple Apparatus for Individual C. elegans Culture. In: Curran SP, editor. Aging: Methods and Protocols [Internet]. New York, NY: Springer US; 2020 [cited 2021 Aug 26]. p. 29–45. (Methods in Molecular Biology). Available from: https://doi.org/10.1007/978-1-0716-0592-9_3

37. Pittman WE, Sinha DB, Zhang WB, Kinser HE, Pincus Z. A simple culture system for long-term imaging of individual C. elegans. Lab Chip. 2017 Nov 7;17(22):3909–20.

38. Yochem J. Nomarski images for learning the anatomy, with tips for mosaic analysis. WormBook [Internet]. 2006 [cited 2022 Jan 27]; Available from: http://www.wormbook.org/chapters/www_nomarskianatomymosaic/nomarskianatomymosaic.html

39. Ronneberger O, Fischer P, Brox T. U-Net: Convolutional Networks for Biomedical Image Segmentation. ArXiv150504597 Cs [Internet]. 2015 May 18 [cited 2022 Jan 28]; Available from: http://arxiv.org/abs/1505.04597

40. Güler RA, Neverova N, Kokkinos I. DensePose: Dense Human Pose Estimation In The Wild. ArXiv180200434 Cs [Internet]. 2018 Feb 1 [cited 2021 Jan 29]; Available from: http://arxiv.org/abs/1802.00434

41. Byerly L, Cassada RC, Russell RL. The life cycle of the nematode Caenorhabditis elegans: I. Wild-type growth and reproduction. Dev Biol. 1976 Jul 1;51(1):23–33.

42. McCulloch D, Gems D. Body size, insulin/IGF signaling and aging in the nematode Caenorhabditis elegans. Exp Gerontol. 2003 Jan 1;38(1):129–36.

43. Fujiwara M, Sengupta P, McIntire SL. Regulation of Body Size and Behavioral State of C. elegans by Sensory Perception and the EGL-4 cGMP-Dependent Protein Kinase. Neuron. 2002 Dec 19;36(6):1091–102.

44. Galimov ER, Pryor RE, Poole SE, Benedetto A, Pincus Z, Gems D. Coupling of Rigor Mortis and Intestinal Necrosis during C. elegans Organismal Death. Cell Rep. 2018 Mar 6;22(10):2730–41.

45. Colaco S, Han DS. Facial Keypoint Detection with Convolutional Neural Networks. In: 2020 International Conference on Artificial Intelligence in Information and Communication (ICAIIC). 2020. p. 671–4.

46. Longpre S, Sohmshetty A. Facial Keypoint Detection. :8.

47. Kinser HE, Mosley MC, Plutzer IB, Pincus Z. Global, cell non-autonomous gene regulation drives individual lifespan among isogenic C. elegans. Gruber J, Tyler JK, editors. eLife. 2021 Feb 1;10:e65026.

48. Sinha DB, Pincus ZS. High temporal resolution measurements of movement reveal novel early-life physiological decline in C. elegans. PLOS ONE. 2022 Feb 2;17(2):e0257591.

49. Fritsch FN, Butland J. A Method for Constructing Local Monotone Piecewise Cubic Interpolants. SIAM J Sci Stat Comput. 1984 Jun 1;5(2):300–4.

50. SciPy 1.0: fundamental algorithms for scientific computing in Python | Nature Methods [Internet]. [cited 2022 Mar 11]. Available from: https://www.nature.com/articles/s41592-019-0686-2

51. Anthony SM, Granick S. Image Analysis with Rapid and Accurate Two-Dimensional Gaussian Fitting. Langmuir. 2009 Jul 21;25(14):8152–60.

